# Cholangiocytes’ Primary Cilia Regulate DNA Damage Response and Repair

**DOI:** 10.1101/2025.01.28.635267

**Authors:** Estanislao Peixoto, Kishor Pant, Seth Richard, Juan E. Abrahante, Wioletta Czaja, Sergio A. Gradilone

## Abstract

Primary cilia have been considered tumor-suppressing organelles in cholangiocarcinoma (CCA), though the mechanisms behind their protective role are not fully understood. This study investigates how the loss of primary cilia affects DNA damage response (DDR) and DNA repair processes in CCA. Human cholangiocyte cell lines were used to examine the colocalization of DNA repair proteins at the cilia and assess the impact of experimental deciliation on DNA repair pathways. Deciliation was induced using shRNA knockdown or CRISPR knockout of IFT20, IFT88, or KIF3A, followed by exposure to the genotoxic agents cisplatin, methyl methanesulfonate (MMS), or irradiation. Cell survival, cell cycle progression, and apoptosis rates were evaluated, and DNA damage was assessed using comet assays and γH2AX quantification. An *in vivo* liver-specific *IFT88* knockout model was generated using Cre/Lox recombination. Results showed that RAD51 localized at the cilia base, while ATR, PARP1, CHK1 and CHK2 were found within the cilia. Deciliated cells displayed dysregulation in critical DNA repair. These cells also showed reduced survival and increased S-phase arrest after genotoxic challenges as compared to ciliated cells. Enhanced DNA damage was observed via increased γH2AX signals and comet assay results. An increase in γH2AX expression was also observed in our *in vivo* model, indicating elevated DNA damage. Additionally, key DDR proteins, such as ATM, p53, and p21, were downregulated in deciliated cells after irradiation. This study underscores the crucial role of primary cilia in regulating DNA repair and suggests that targeting cilia-related mechanisms could present a novel therapeutic approach for CCA.

New and Noteworthy: Our findings reveal a novel connection between primary cilia and DNA repair in cholangiocytes. We showed that DDR and DNA repair proteins localize to cilia, and that deciliation leads to impaired cell survival and S-phase arrest under genotoxic stress. Deciliated cells exhibit heightened DNA damage, evidenced by increased γH2AX signals and comet assay results, a phenotype mirrored in *in vivo* IFT88 knockout mice. Furthermore, key DDR regulators, including ATM, p53, and p21, are downregulated in deciliated cells following irradiation, highlighting a crucial role for primary cilia in maintaining genome stability.

## INTRODUCTION

Cholangiocarcinoma (CCA), also known as bile duct cancer, represents a particularly lethal malignancy within the spectrum of liver cancers. Characterized by the malignant transformation of cholangiocytes—the epithelial cells lining the bile ducts in the liver—CCA poses significant treatment challenges due to typically late diagnoses and limited therapeutic options (1–4). Despite recent advancements in understanding liver cancers, CCA remains difficult to manage, with pharmacological treatments often showing inadequate responses and surgical intervention viable only in the early stages (5). Consequently, most patients face a dismal median survival time of less than two years, underscoring the urgent need for novel therapeutic strategies (5).

Primary cilia are multisensory organelles ubiquitously expressed in epithelial cells that act as cellular antennae to detect signals from the extracellular environment (6, 7). While cholangiocytes typically exhibit primary cilia, we and others have observed that their expression is notably diminished in CCA cells (8–10). The experimental removal of cilia through methods such as the deletion of *IFT88*, a gene required for ciliogenesis, results in a shift towards malignant-like behavior characterized by increased proliferation and the induction of anchorage-independent growth. Furthermore, the restoration of cilia on malignant cells reverses these effects, suggesting a tumor-suppressing role for these organelles (8). Previous results revealed the involvement of histone deacetylase 6 (HDAC6) and Sirtuin-1 in the loss of cilia within CCA cells, while inhibition of either enzyme leads to the re-expression of cilia (8, 11, 12). The precise mechanisms through which primary cilia exert this protective influence against cancer development remain to be fully elucidated.

DNA repair processes critical for carcinogenesis suppression appear intimately associated with primary cilia, especially within the centrosome at the cilia’s basal body. Disrupting ciliogenesis affects the DNA damage response, as evidenced by altered DNA-PK activity in conditions like glioblastoma (13, 14). Key DNA repair and damage response proteins, including BRCA1, BRCA2, PARP1, NBS1, ATM, ATR, CHK1, CHK2, and p53 are associated with this structure (15–17). CEP164, a centrosomal protein crucial for ciliary formation, interacts with both ATR (Ataxia Telangiectasia and Rad3-related protein) and ATM (Ataxia Telangiectasia Mutated protein). CEP164 undergoes phosphorylation in response to DNA damage, and its resulting downregulation leads to reduced phosphorylation of other proteins in the DNA damage response, such as CHK2 and CHK1 (18). Proteins linked to ciliopathies, such as MRE11, ZNF423, and CEP164, play roles in the DNA damage response (19). Furthermore, NEK proteins participate in primary cilia function and DNA damage response (20). NEK8/NPHP9 is a ciliary kinase associated with the ciliopathies nephronophthisis and polycystic kidney disease. NEK8 suppresses double-strand breaks (DSB) by limiting cyclin A-associated CDK activity, and NEK8 mutant mice accumulate DNA damage (21).

The presence of multiple DNA repair proteins at the ciliary basal body and the alterations observed in cells lacking cilia suggest that the loss of primary cilia in cholangiocytes might significantly affect DNA repair rates. If so, the absence of primary cilia during cholangiocarcinogenesis would induce compromised DNA repair mechanisms. To test this hypothesis, we employed various genotoxic agents targeting different DNA repair pathways and minimized protein-specific biases using different CRISPR knockouts of key ciliary proteins. Our results from in vitro experiments and an in vivo Alb-Cre IFT88 mouse model indicate increased DNA damage in cilia-deficient cells, impacting both the cellular and molecular levels and potentially contributing to tumor progression. These findings may offer new insights into therapeutic strategies for restoring genome stability in cholangiocarcinogenesis.

## MATERIALS AND METHODS

### Cell Lines and Culture

Normal human cholangiocyte (NHC) cells were cultured in Dulbecco’s Modified Eagle Medium (DMEM, Sigma-Aldrich, St. Louis, MO, USA) supplemented with 10% fetal bovine serum (FBS), 100 U/ml penicillin, 100 µg/ml streptomycin, 25 µg/ml adenine, 1 µg/ml epinephrine, 5 µg/ml insulin, 13.6 ng/ml Triiodo-L-thyronine, 8.3 µg/ml holo-transferrin, 1.3 µg/ml hydrocortisone, 10 mg/ml gentamicin, and 10 ng/ml epidermal growth factor (EGF). The HuCCT1 cell line was grown in Ham’s F-12 Medium (F-12) supplemented with 10% FBS, 100 U/ml penicillin, 100 µg/ml streptomycin, and 10 mg/ml gentamicin. Cultures were maintained at 37°C in a humidified atmosphere with 5% CO2. Cells were exposed to irradiation using the Rad Source RS 2000 Biological Research Irradiator, which generates X-ray ionizing radiation fields, treated with methyl methanesulfonate (MMS, Sigma-Aldrich) or with cisplatin (Tocris, Minneapolis, MN, USA).

### shRNA Transfection and CRISPR-Cas9 gene editing

Cells were stably transfected with lentiviral-mediated shRNA vectors to knock down IFT88. CRISPR-Cas9 [clustered regularly interspaced short palindromic repeats (CRISPR)–CRISPR-associated enzyme 9] was used to knockout ciliary proteins, including kinesin family (KIF)-3A (gRNA 5’-AGAAAGCUGCGAUAAUGUGA-3’), IFT20 (gRNA 5’-GAGCAAGUUCCGAGCACCGA-3’) and IFT88 (gRNA 5’-UUCAUUAACUGAAUACUGAC-3’) in NHC cells (Supp. Fig. 1). NT-shRNA and NT-gRNA were used as controls.

### RNA-Seq Analysis

Quality control, data alignment and gene quantification were analyzed using the CHURP pipeline (22) at the University of Minnesota Supercomputing Institute (MSI). 2 x 75bp FASTQ paired-end reads for 12 samples (31.02 million reads average per sample) were trimmed using Trimmomatic (v0.33) enabled with the optional “-q” option; 3bp sliding-window trimming from 3’ end requiring minimum Q30. Quality control on raw sequence data for each sample was performed with FastQC. Read mapping was performed via HISAT2(v2.1.0) using the human genome (GRCh38.97) as reference. Gene quantification was done via Feature Counts for raw read counts. Differentially expressed genes were identified using the edgeR (negative binomial, R programming) feature in CLCGWB (Qiagen, Redwood City, CA) using raw read counts. We filtered the generated list based on a minimum 2X absolute fold change and FDR corrected p < 0.05. Differential pathway analysis was performed by ReactomeGSAing. using correlation adjusted mean rank gene set test (Camera) algorithm. Differentially expressed pathways with a p-value below 0.05 were considered significant. Raw RNAseq data were deposited on the NCBI Gene Expression Omnibus (GEO) repository under the accession number GSE280249.

### Survival and Cell Proliferation Assay

Cell proliferation was assessed using live-cell imaging assays. Specifically, 5×10^3^ cells were seeded in 100 μL of culture media per well in 96-well plates. These were then incubated at 37°C and 5% CO_2_, and the IncuCyte live-cell analysis system (Essen BioScience, Ann Arbor, MI, USA) was used for continuous monitoring. Data were processed and analyzed using IncuCyte S3 software.

### Analysis of Cell Cycle

Cells were harvested with 0.025% trypsin + 5 mM EDTA in DPBS, with 2.5% FBS + 5 mM EDTA added when cells detached. Cells were washed with cold DPBS and fixed overnight at – 20°C with 70% ethanol and 30% DPBS. Fixed cells were washed with DPBS and incubated with 20ug/ml propidium iodide and 200 ug/ml RNase A for 30 minutes at room temperature in the dark. Cells were analyzed on a Becton Dickinson Fortessa X-20 flow cytometer (BD Biosciences, San Jose, CA, USA). Intact cells were gated in the FSC/SSC plot to exclude small debris and on a PI plot of PI-A vs. PI-W to exclude doublets. Cell cycle was determined using ModFit LT software version 5.0 (Verity Software House, Inc., Topsham, ME).

### Apoptosis Assay

To assess apoptosis, cells were detached using 0.025% trypsin supplemented with 5 mM EDTA in PBS. To neutralize the trypsin, 2.5% FBS in PBS with an additional 5mM EDTA was added immediately upon cell detachment. The cell suspension was centrifuged, and the pellets were washed with cold PBS, and subsequently stained with Annexin V-FITC and propidium iodide (PI) for 15 minutes at room temperature in the dark. Flow cytometry was performed as described above. Apoptotic cells were identified based on their staining patterns: cells in early apoptosis appeared in the lower right quadrant of the Annexin V-FITC/PI dot plot (indicating Annexin V-FITC staining only), while cells in late apoptosis or necrosis were located in the upper right quadrant (showing both Annexin V-FITC and PI staining). Data were analyzed using BD FACS DIVA software version 9.0.

### Comet Assay for DNA Damage Evaluation

DNA damage was assessed using the OxiSelect Comet Assay Kit (Cell Biolabs Inc., San Diego, CA, USA), with slight modifications to the standard protocol. After being treated with genotoxic agents for specified durations, NHC cells were collected, washed twice in PBS, and resuspended in 75 μL of OxiSelect Comet Agarose (prepared in PBS at pH 7.4 and maintained at 37°C). This cell-agarose mixture was promptly transferred onto glass microscope slides pre-coated with 1% (w/v) normal melting point agarose and allowed to solidify for 10 minutes at 4°C. Subsequently, to facilitate protein removal, the slides were submerged in lysis solution overnight at 4°C, followed by a 4-hour exposure to alkaline solution (0.3 M NaOH and 1 mM EDTA). Electrophoresis was then conducted in an alkaline electrophoresis solution containing 0.3 M NaOH and 1 mM EDTA at 22 V for 35 minutes. Post-electrophoresis, slides were washed thrice with chilled distilled water (2 minutes each) and once with 70% ethanol for 5 minutes. The DNA was stained with 40 μL of Vista Green DNA Dye for visualization. The stained slides were examined under a Zeiss Axio Observer microscope equipped with Apotome, facilitating detailed analysis of the DNA damage manifested as comet tails. Comet Score (v2.0.0.0) was used to quantify the Olive tail moment.

### Immunofluorescence Assay

Cells were plated on four-well chamber slides and allowed to grow for one day. Following treatments with genotoxic agents, the cells underwent a double wash with PBS and were then fixed with ice-cold methanol for 5 minutes, followed by three washes in 0.1% PBS with Tween 20 (PBST). To block non-specific binding, cells were incubated in 3% bovine serum albumin (BSA) in PBS for 1 hour at room temperature. Subsequently, they were incubated overnight at 4°C with the following primary antibodies diluted in PBS containing 1% BSA and 0.1% Tween: mouse anti-gamma tubulin (1:200, Sigma-Aldrich #T5326), rabbit or mouse anti-ARL13B (1:200, Proteintech, Rosemont, IL, USA #17711-1-AP and #66739-1-Ig respectively), rabbit anti-γH2AX (1:400, Cell Signaling Technology, Danvers, MA, USA #9718S), rabbit anti-PARP1 (1:200, Proteintech, 13371-1-AP), rabbit anti-human RAD51 (1:200, Proteintech #14961-1-AP), mouse anti-human ATR (1:200, Santa Cruz Biotechnology, Dallas, TX, USA #sc-515173), rabbit anti-human CHK1 (polyclonal, 1:200, Proteintech #25887-1-AP), or mouse anti-CHK2 (1:200, Cell Signalling #3440S). After rinsing thrice with PBST, cells were incubated with Alexa Fluor 594-conjugated goat anti-rabbit and/or Alexa Fluor 488-conjugated goat anti-mouse secondary antibodies (1:200, Life Technologies, CA, USA) for 1 hour, followed by three additional PBST washes. Nuclei were stained with DAPI using ProLong Gold Antifade Reagent (Invitrogen). Immunofluorescence images were acquired and analyzed using a Nikon D-eclipse C1 Si microscope or a Zeiss Axio Observer with Apotome. The number of foci per nuclei was counted using CellProfiler.

### Western Blot

Cells were rinsed with ice-cold PBS and lysed using RIPA buffer containing 50 mM Tris-HCl (pH 7.5), 0.1% sodium dodecyl sulfate (SDS), 0.1% Triton X-100, 1% Nonidet P-40, 0.5% sodium deoxycholate, 150 mM NaCl, and 1 mM phenylmethylsulfonyl fluoride. Lysates were incubated overnight, then centrifuged at 13,000 rpm for 15 minutes at 4°C to pellet insoluble material. Protein concentrations were quantified using the Pierce BCA Protein Assay Kit (Thermo Scientific, Waltham, MA, USA). Proteins were separated on a 4-20% SDS-polyacrylamide gel and transferred onto nitrocellulose membranes (Bio-Rad Laboratories, Hercules, CA, USA). Membranes were blocked with 5% skim milk in TBST for 1 hour at room temperature, washed with TBST, and incubated overnight at 4°C with primary antibodies diluted in TBST/5% BSA directed against rabbit anti-actin (1:1000, Sigma-Aldrich #A2066), rabbit anti-β-Actin (1:1000, Cell Signaling Technology, 4970S), rabbit anti-p21 Waf1/Cip1 (1:1000, Cell Signaling Technology #2947), mouse anti-p53 (1:1000, Santa Cruz Biotechnology #sc-126), and mouse anti-ATM (1:1000, Santa Cruz Biotechnology #sc-377293). Following three TBST washes, membranes were incubated with HRP-conjugated secondary antibodies in 5% skim milk in TBST for 1 hour at room temperature. Detection was performed using the Pierce ECL Western Blotting Substrate (Thermo Scientific, Rockford, IL, USA).

### Animal studies

Alb-Cre-IFT88KO mice on a C57BL/6J background were housed in a dedicated animal facility under a 12-hour light/dark cycle with ad libitum access to a standard diet and water. Liver samples were obtained from 48-week-old mice. All animal procedures were conducted in accordance with protocols approved by the University of Minnesota Institutional Animal Care and Use Committee (IACUC).

### Statistical analysis

Data are presented as mean ± standard error (SE). Statistical comparisons were conducted using one-or two-way ANOVA, Student’s unpaired t-test or the Mann-Whitney U test, as appropriate. Significance levels were set at P < 0.05, P < 0.01, P < 0.001, and P < 0.0001. Experiments were conducted at least twice when using different clones of the same mutation, as well as different CRISPR mutations targeted for ciliary loss. If only a single clone was used, experiments were performed independently at least three times.

## RESULTS

### DNA damage response proteins colocalize with ciliary structures

We first asked if DNA damage response (DDR) proteins and DNA repair proteins colocalize at the base of primary cilia in cholangiocytes. Normal Human Cholangiocyte (NHC) cells are well-established and widely used cholangiocytes derived from nondiseased patient tissue (23). Immunofluorescence assays in NHC cells revealed that RAD51 localizes at the base of the cilia. Additionally, ATR, PARP1, CHK1 and CHK2 exhibited colocalization within the ciliary structure (Figure 1A). This suggests a spatial association between these DDR proteins and the primary cilium, consistent with a potential functional interplay in DNA repair processes.

**Figure 1.**
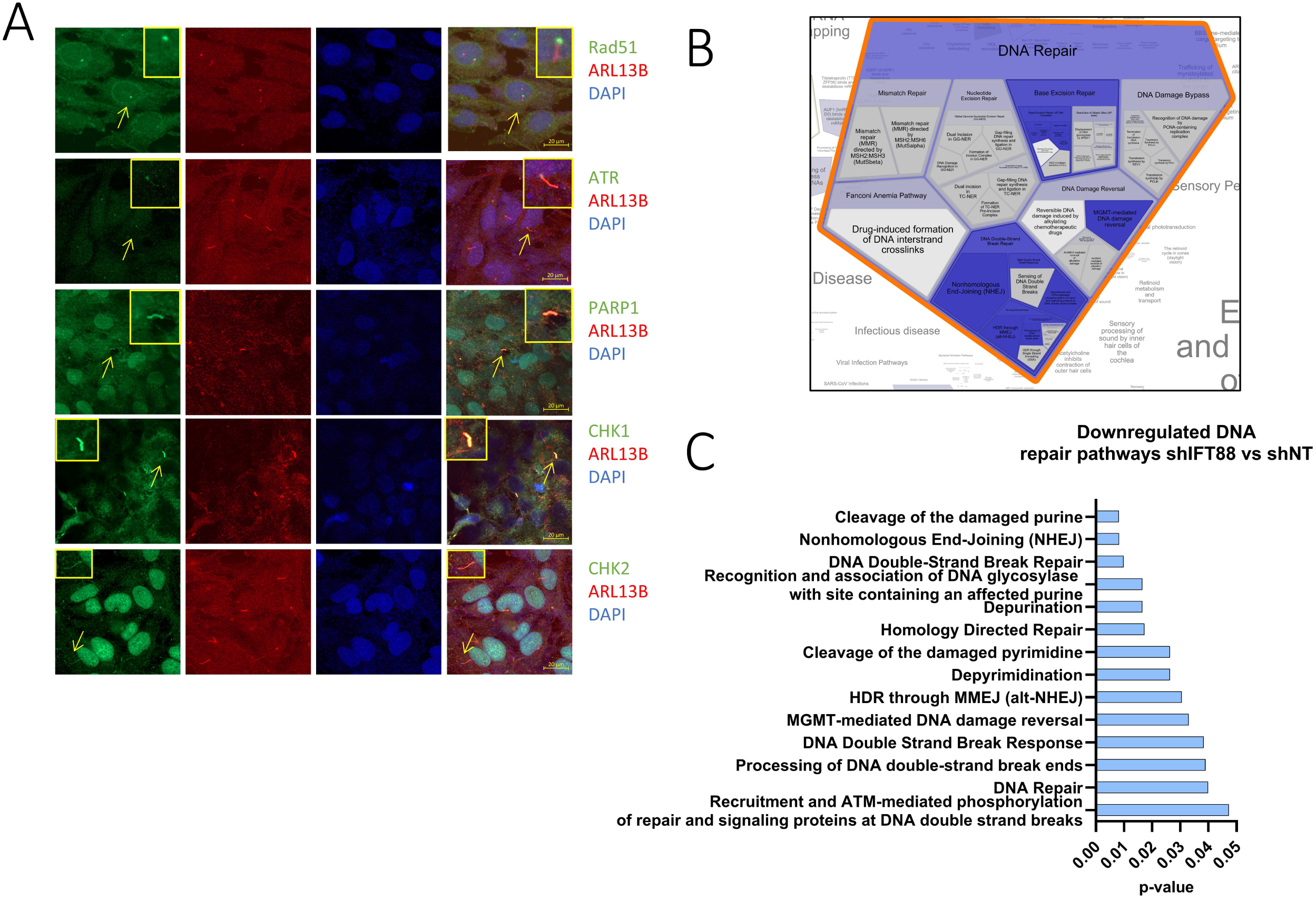
DNA damage response genes colocalize with the base of the cilia and the deciliation process impacts DDR and DNA repair pathways. (A) Immunofluorescence micrographs of RAD51, ATR, CHK1, CHK2, PARP1 and the ciliary marker ARL13B in NHC cells. 630X, insets show a magnification of ciliary structures indicated by the yellow arrow. (B) Voronoi diagram representation of the ‘DNA repair’ pathway from ReactomeGSA pathway enrichment analysis, using all expressed mRNA elements from NHC clonal shRNA shNT and shIFT88 cells. Blue indicates pathways downregulated in shIFT88 cells. (C) Downregulated DNA repair pathways identified by ReactomeGSA in shIFT88 cells compared to shNT cells, with their corresponding p-values.

To further explore the relationship between DDR mechanisms and primary cilia, we conducted RNA sequencing and pathway analysis of NHC cells with and without cilia. Cells with a knockout or knockdown of IFT88, KIF3A, or IFT20 cannot express primary cilia (8, 24, 25), and are referred to as ‘deciliated’. Cells were stably transfected with lentiviral-mediated shRNA vectors to knock down IFT88 and induce deciliation; these lines are referred to as shIFT88. Pathway analysis performed using ReactomeGSA (26, 27) revealed downregulation of pathways associated with base excision repair, DNA damage reversal, and DNA double-strand break repair in deciliated NHC cells (Figure 1B, C)). Our findings show that key DDR and repair proteins localize to ciliary structures in cholangiocytes, suggesting a spatial and functional link between primary cilia and DDR mechanisms. Furthermore, the pathway analysis results suggest that primary cilia may contribute to genome stability and support further investigation.

### The loss of primary cilia impairs cholangiocyte survival under genotoxic stress

If deciliated cells have impaired DNA damage response and repair pathways, they should exhibit reduced survival when challenged with genotoxins compared to ciliated cells. To investigate this hypothesis, we generated deciliated NHC clones by knocking out key ciliary assembly proteins (KIF3A, IFT88, or IFT20, Fig. Supp. 1) using CRISPR-Cas9 and assessed the proliferation rates of normal (CRISPR non-targeting control NT) and deciliated NHC clones using live-cell imaging. At each hourly time point, proliferation was normalized to the corresponding untreated controls using the formula: (area confluence of treated clone × 100) / area confluence of untreated clone. We employed various genotoxic approaches, including irradiation to induce primarily double-strand breaks (28), methyl methanesulfonate (MMS) to cause single-strand breaks (29), and cisplatin to induce DNA crosslinking (30). Ciliated and deciliated cells were treated with different genotoxins and immediately placed under Incucyte monitoring. We found a significant reduction in the survival of deciliated NHC cells 72 hours after a 1-hour treatment with 1 mM methyl methanesulfonate, a 24-hour treatment with 2.5 µM cisplatin, or exposure to 5 Gy of irradiation, in comparison to their control ciliated counterparts (Figure 2A). Some clones did not show decreased survival, which may be due to variability among CRISPR clones caused by off-target edits, epigenetic modifications, or genetic drift. To account for this variability, we maximized the use of different clones in our analysis.

**Figure 2.**
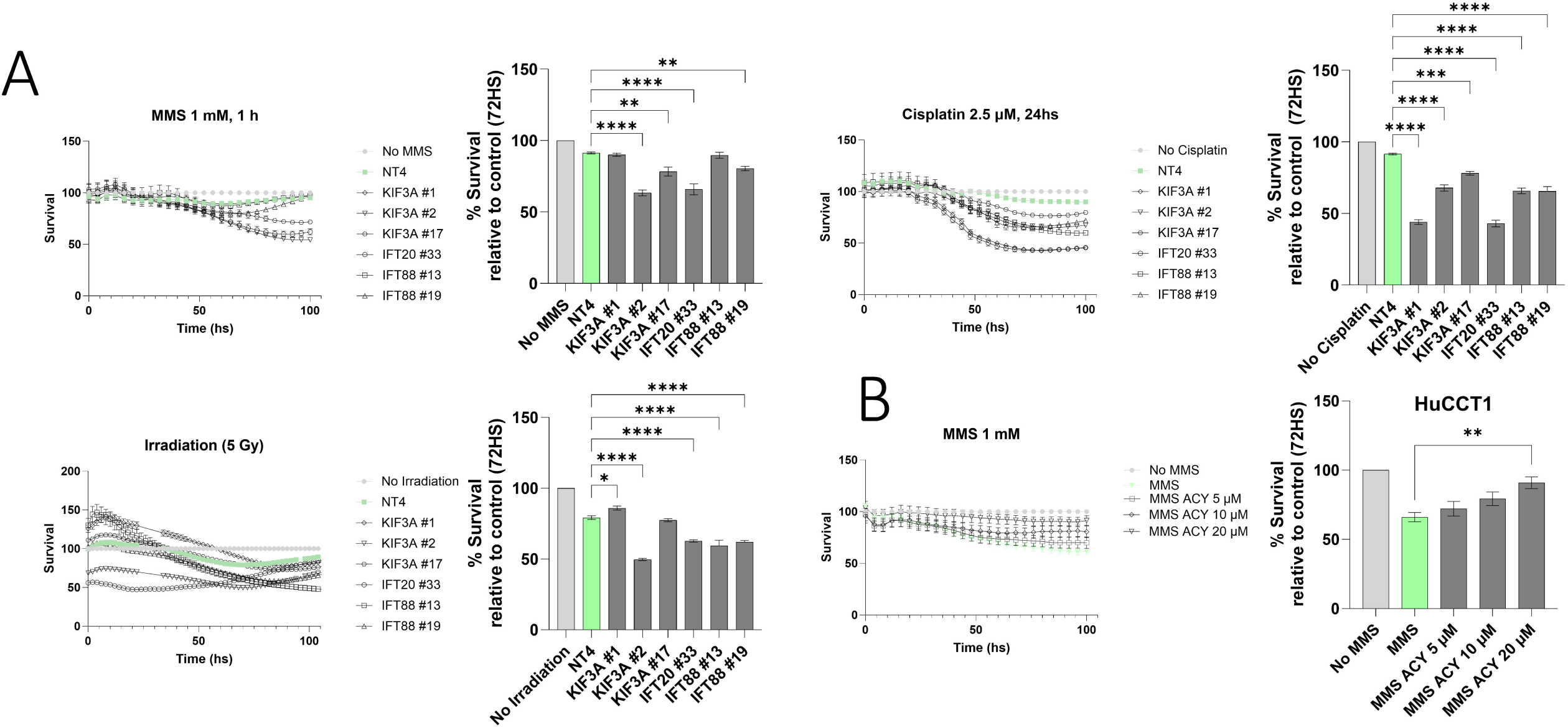
Deciliated cholangiocytes show lower survival than normal cholangiocytes when challenged by genotoxins. (A) Survival curves of cilated and deciliated clonal CRISPR knockout NHC cell lines treated with 1 mM MMS, 2.5 µM cisplatin or 5Gy relative to their respective controls (One-way ANOVA, *p<0.05, **p<0.01, ***p<0.001, ****p<0.0001). (B) Survival curves of CCA cells treated with ACY1215 (5, 10 or 20 µM) or MMS (1 mM, One-way ANOVA, **p<0.01.

Tumoral cells such as HuCCT1, which do not express cilia, can recover ciliary expression by treating them with an HDAC6 inhibitor like ACY1215 (ACY) (11). To provide additional evidence that cilia promote cellular survival under genotoxic stress, we challenged ACY-induced ciliogenesis HuCCT1 cells with MMS at different ACY concentrations and monitored survival. Survival was calculated as: area confluence of HuCCT1 cells treated with [ACY] + MMS × 100) / area confluence of HuCCT1 [ACY] without MMS. An increase in survival was observed in ACY-treated cells as the ACY concentration increased, with a statistically significant effect at 20 µM compared to untreated control tumor cells following MMS exposure. (Figure 2B). Taken together, these data show that the presence of primary cilia enhances cellular survival during genotoxic stress.

To further investigate the impact of cilia loss on cell survival mechanisms, we analyzed the cell cycle distribution and apoptosis rates in genotoxically challenged cells. We exposed ciliated and deciliated cells to cisplatin for 24 hours, followed by cell cycle analysis. Additionally, cells were treated with MMS for 1 hour or irradiated with 5 Gy, and cell cycle analysis was performed 24 hours later. MMS-challenged cells lacking cilia demonstrated a statistically significant increase in the percentage of cells found in S phase as compared to cells with cilia (Figure 3, Fig. Supp. 2). This may suggest that stalled replication due to DNA damage actively prevents progression through the cell cycle in the absence of primary cilia. For irradiated deciliated cells, we observed a slight, non-significant increase in S phase, except in KIF3A mutants. In response to cisplatin treatment, we noted a slight non-significant trend decrease in G2 phase arrest in deciliated cells, except in IFT88-deficient cells. We also assessed apoptosis rates in ciliated and decilated cells that had been exposed to genotoxic stress. Compared to their ciliated counterparts, deciliated irradiated cells exhibited a slight increase in apoptosis rates (Fig. Supp. 2). This elevation in apoptosis suggests that the inability of deciliated cells to effectively repair DNA damage may contribute to their increased susceptibility to programmed cell death, though more research is needed. Taken together, our results indicate that the lack of cilia exacerbates the effects of genotoxins on cell survival and arrest. This effect appears to be largely due to stalled replication caused by DNA damage, which could actively prevent cell cycle progression in the absence of primary cilia.

**Figure 3.**
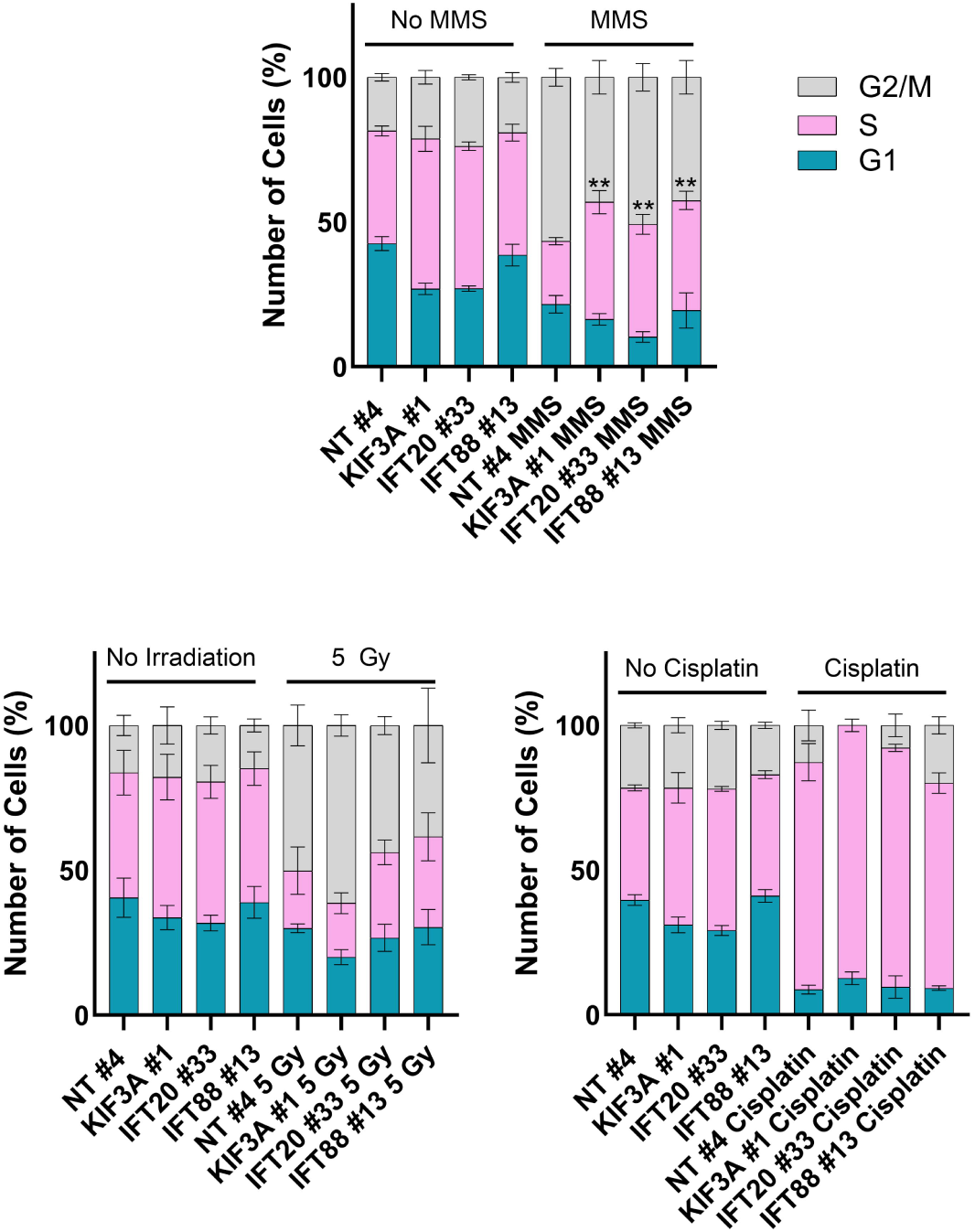
Cell cycle dynamics of ciliated and deciliated clonal CRISPR knockout cell lines in response to genotoxic treatments. Stacked bar graphs representing the percentage of cells in each step of the cell cycle of ciliated and deciliated cells on basal conditions or 24 hrs after 2.5 µM cisplatin, 1 mM MMS or 5 Gy. One-way ANOVA, **p<0.01 for the S-phase comparison between NT #4 MMS-treated cells and MMS-treated deciliated clones.

### Deciliated cholangiocytes show increased DNA damage

Phosphorylated H2AX (γH2AX) is an established marker of DNA double-strand breaks and plays a crucial role in recruiting DNA repair machinery to sites of damage. Its absence can lead to genomic instability and contribute to cancer progression (31). We investigated whether the absence of cilia in genotoxin-challenged cells alters H2AX levels. To address this, we performed immunofluorescence on ciliated and deciliated cells treated with cisplatin for 24 hours, irradiated with 5 Gy, or exposed to 1 mM MMS, and evaluated them 24 hours later. We observed elevated γH2AX levels in deciliated normal human cholangiocyte (NHC) cells exposed to 5 Gy of radiation at 24 hours post-exposure (Fig 4A). Furthermore, RAD51, essential for homologous recombination (HR) repair of DSBs and homologous recombination marker (32), increased levels in deciliated NHC cells 6 and 24 hours after 5 Gy radiation exposure compared to irradiated controls, hinting at RAD51 activity dysregulation (Figure Supp. 3). Additionally, deciliated NHC cells treated with 2.5 mM cisplatin for 24 hours or with MMS for 1 hour, followed by a 24-hour recovery, exhibited higher γH2AX levels compared to ciliated controls. (Figure 4B, C). Taken together, these results suggest either enhanced γH2AX activity or impaired resolution of DNA damage in deciliated cells.

**Figure 4.**
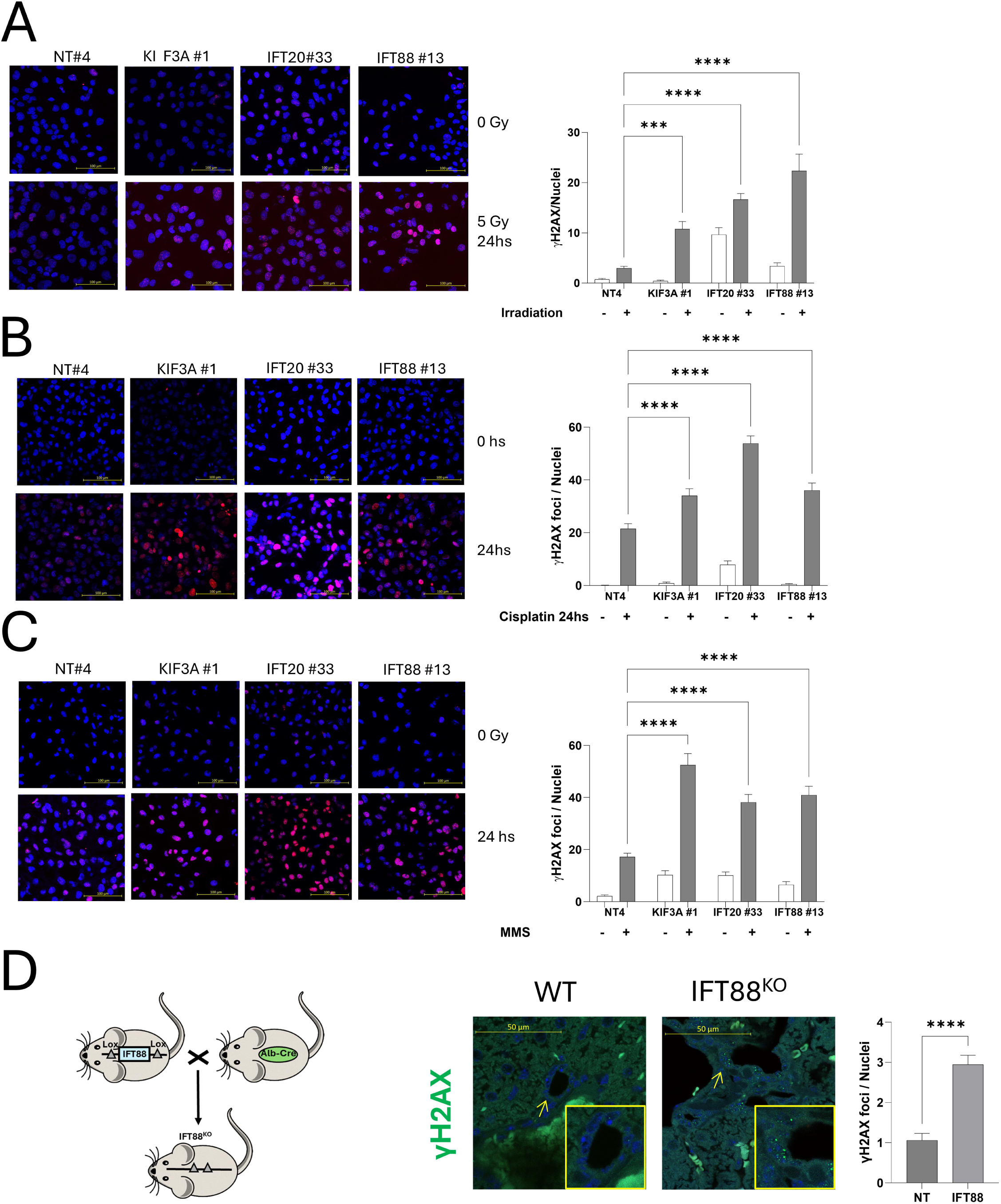
γH2AX is increased on deciliated clonal CRISPR knockout cell lines after genotoxins. (A) Immunofluorescence analysis of γH2AX in ciliated and deciliated NHC cells 24 hours after 5 Gy irradiation or in untreated controls. (One Way Anova, ****p<0.0001, 630X). (B) Immunofluorescence for γH2AX of ciliated and deciliated NHC cells at 24 hrs after treatment with 2.5 µM cisplatin (One-way ANOVA, ****p<0.0001, 630X). (C) Immunofluorescence for γH2AX of ciliated and deciliated NHC cells at 24 hrs after treatment with 1 mM MMS. One-way ANOVA, ****p<0.0001, 630X. (D) Immunofluorescence staining and quantification of γH2AX in IFT88 knockout mice (n=3, unpaired t-test, ****p<0.0001, magnification x630). The inset shows the IFT88/LOX construct, illustrating the Albumin-Cre–mediated conditional knockout design.

Since our *in vitro* studies indicated increased DNA damage in cells lacking primary cilia, we investigated whether a similar effect occurs *in vivo* using IFT88 knockout mice. Since Cre is driven by an albumin promoter, it facilitates the recombination of loxP sites in hepatocytes and in the liver precursor cells of hepatocytes and cholangiocytes during embryonic development (33). This model has also been used in previous studies to induce cholangiocarcinoma (34–36). As expected, Alb-Cre-IFT88 knockout mice displayed a significant reduction in primary cilia expression on cholangiocytes (37) and is also known to induce biliary hyperplasia (38, 39). We performed γH2AX immunofluorescence on liver samples from these mice and found that 48-week-old IFT88 KO animals exhibited elevated γH2AX levels in bile duct cells compared to controls (Figure 4D). This suggests that the absence of primary cilia may make cells more vulnerable to physiological stress, including genotoxic agents derived from hepatic metabolism (40).

γH2AX foci may increase due to enhanced DNA repair activity or an accumulation of unresolved DNA damage. To further quantify the effect of cilia on DNA damage, we performed a comet assay (single-cell gel electrophoresis) under alkaline conditions, which detects DNA fragments corresponding to single– and double-strand breaks, using NHC cells challenged with 1 mM MMS for 1 hour or 10 Gy irradiation as genotoxic agents. Cisplatin forms DNA crosslinks (41), which hinder DNA fragment migration; therefore, it was excluded from these experiments The majority of deciliated cell clones exhibited a significantly increased Olive moment (% of DNA within the tail multiplied by the length between the centers of the head and tail), an indicator of DNA damage, compared to cells with intact cilia 24 hours after treatment with MMS or exposure to irradiation (Figure 5A, B), supporting the hypothesis that they accumulate unresolved DNA damage.

**Figure 5.**
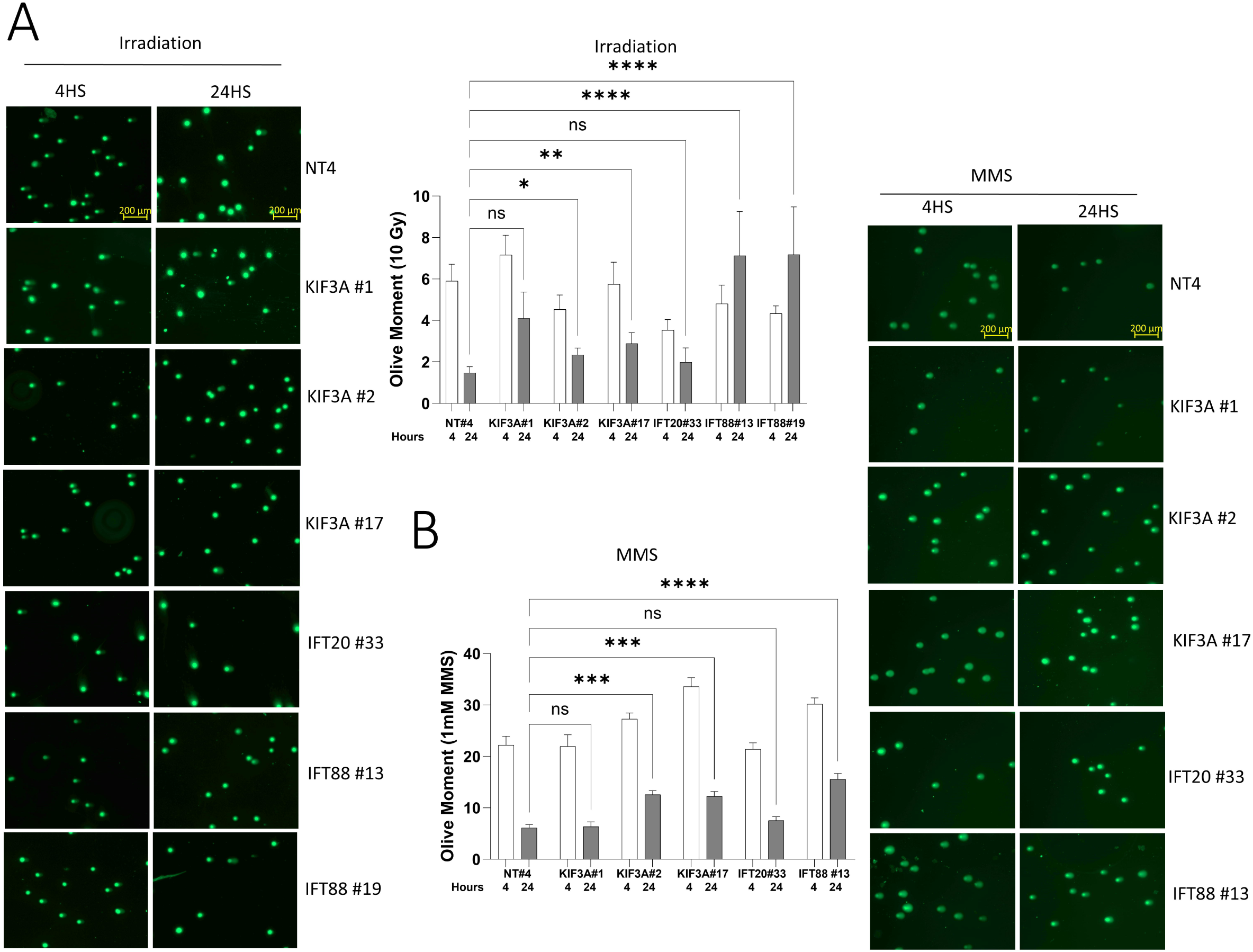
DNA damage is increased in deciliated clonal CRISPR knockout cell lines after exposure to genotoxins. Representative pictures and comet assays of ciliated and deciliated NHC cells (clones KIF3A #1, KIF3A #2, KIF3A #17, IFT20 #33, IFT88 #13 and IFT88 #19) after 4 or 24 h of treatment with 10 Gy irradiation (A) or 1 mM MMS (B) One-way ANOVA, ***p<0.001, ****p<0.0001, 100X.

### Dysregulation of DNA Damage Response Proteins in Deciliated Cells

In normal ciliated cells, irradiation induces DNA double-strand breaks, leading to the rapid activation of the ATM kinase. ATM phosphorylates and stabilizes p53, a key tumor suppressor, which then drives the expression of downstream targets such as p21. The activation of this pathway in response to DNA damage is crucial for maintaining genomic stability, as it prevents cells with damaged DNA from progressing through the cell cycle. By studying deciliated IFT88 cells, we aimed to determine whether the absence of primary cilia disrupts this ATM-p53-p21 axis and compromises the cell’s ability to manage DNA damage, potentially leading to impaired repair mechanisms or altered cell fate decisions, such as increased apoptosis or genomic instability.

To investigate the DNA damage response, we irradiated ciliated and deciliated cells with 5 Gy, collected protein samples at 30 minutes, 6 hours, and 24 hours, and performed Western blot analysis to examine the expression of ATM, p53, and p21. Deciliated NHC cells exhibited a reduction in ATM levels 24 hours post-5 Gy irradiation (Figure 6A, B). Additionally, deciliated NHC cells showed a decrease in p53 levels at 6 hours following 5 Gy irradiation (Figure 6C). Moreover, we observed an increase in p21 levels in NHC cells 24 hours after irradiation (Figure 6D) but the increase was lower on deciliated cells. These findings indicate dysregulation of key DNA damage response proteins in deciliated cells.

**Figure 6.**
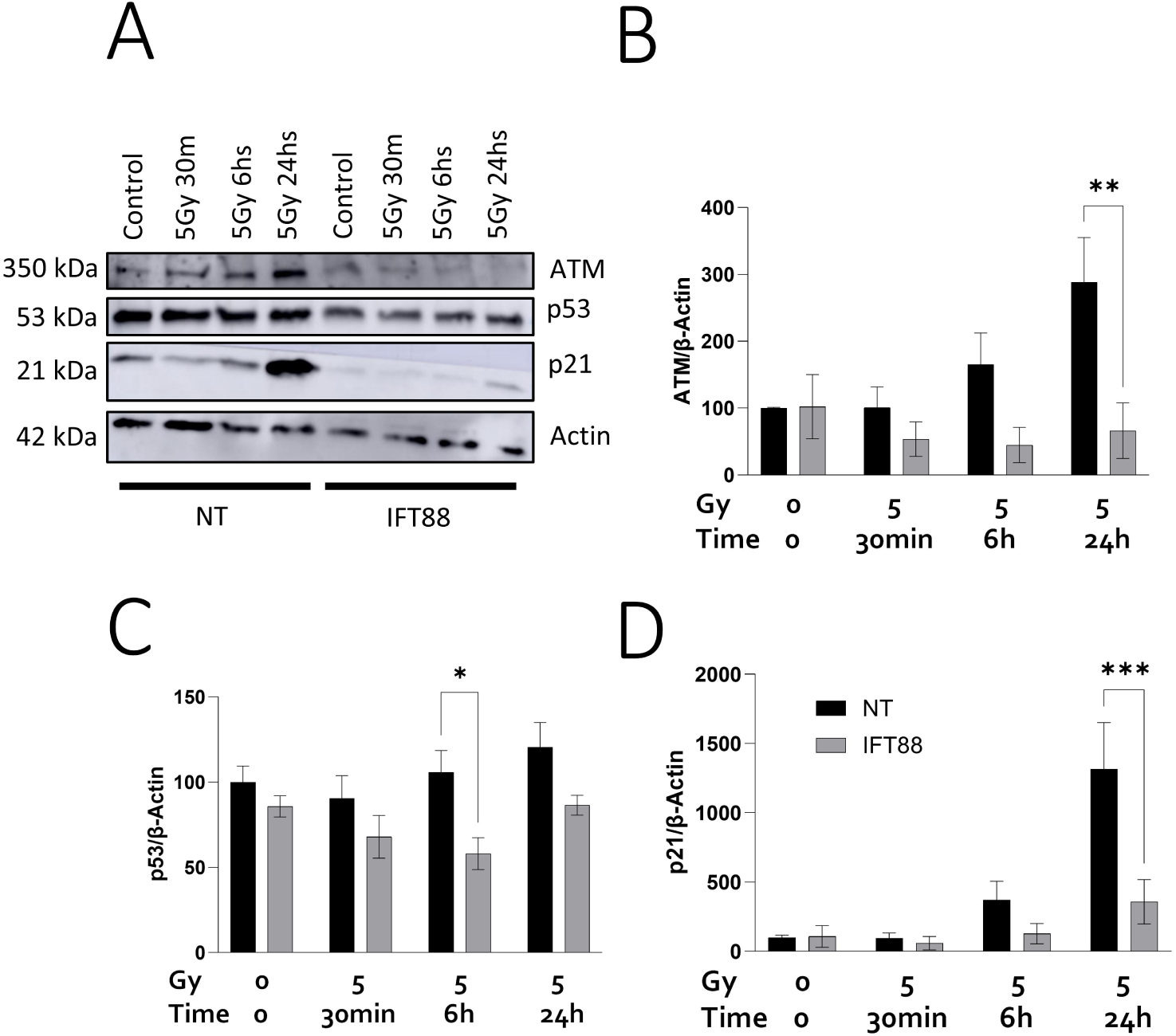
DNA damage response proteins are dysregulated on deciliated clonal CRISPR knockout cell lines after irradiation. (A) Western blot analysis of control and IFT88-deficient cells following 5 Gy irradiation, using antibodies against ATM, p53, p21, and actin. (B) ATM Western blot analysis of ciliated and deciliated NHC cells treated with 5 Gy (2-way ANOVA, **p<0.01, ***p<0.001). (C) p53 Western blot analysis of ciliated and deciliated NHC cells treated with 5 Gy (2-way ANOVA, *p<0.05, **p<0.01). (D) p21 Western blot analysis of ciliated and deciliated NHC cells treated with 5 Gy. 2-way ANOVA, *p<0.05, **p<0.01.

## DISCUSSION

Previous work has suggested that the lack or impairment of primary cilia, as well as disturbances in the signaling pathways associated to cilia, could facilitate oncogenic processes in cholangiocytes (8, 11, 12, 42). Here, we provide evidence that one contributor to these processes could be DNA repair. Here we show that: i) DDR proteins including Rad51, ATR, PARP1, CHK1, and CHK2 co-localize with ciliary structures within cholangiocytes; ii) the absence of cilia in cholangiocytes leads to an aberrant expression profile of DDR genes; iii) cholangiocytes that are devoid of cilia demonstrate an increased sensitivity towards genotoxic agents; iv) such deciliated cells manifest an elevated level of γH2AX in response to genotoxic stress, while a liver-specific *IFT88* knockout mouse model shows elevated levels of γH2AX in the ciliary defective cholangiocytes; v) an enhanced accumulation of DNA damage is observed in deciliated cells; and vi) exposure to genotoxic stimuli reduces the presence of DDR proteins in deciliated cells. Collectively, these findings underscore, for the first time, the critical role of cilia in maintaining genomic stability in cholangiocytes.

We observed the impact of ciliary loss on DNA repair across three distinct DNA damage models, including alkylations (MMS), double-strand breaks (irradiation), and crosslinks (cisplatin). These results reinforce the idea that primary cilia play a conserved role in orchestrating the DNA damage response. Furthermore, the results were consistently replicated across different genetic approaches to deciliation, including CRISPR knockouts of IFT88, KIF3A, or IFT20, indicating that the process is cilia-dependent rather than protein-specific.

Poly (ADP-ribose) polymerase (PARP) was observed to co-localize with centrosomes in mouse embryonic fibroblasts, human osteogenic sarcoma, and human glioma cells (43, 44). Similarly, ATM, ATR, CHK1, and CHK2 showed co-localization with centrosomes in HeLa cells (45). Our investigations into cholangiocytes revealed a comparable co-localization pattern during interphase, with the proteins ATM, PARP1, CHK1 and CHK2 also present on the cilia. This suggests that the microtubules organized by the centrosomes may serve as a conduit for the trafficking of DNA repair proteins, potentially influenced by ciliary signaling mechanisms. Notably, the efficacy of DNA-damaging agents is augmented by microtubule targeting agents, which disrupt the trafficking pathway of DNA repair proteins (46).

Our results align with and build upon previous findings from different models. Knockout of IFT88 and CEP164 attenuates the DNA damage response via diminished DNA-PK-p53 activation in human retinal pigment epithelial (RPE1) cells (13). Additionally, primary cilia have been implicated in meiotic recombination and the repair of double strand breaks during gametogenesis in zebrafish (47). Moreover, the inhibition of ciliogenesis has been shown to augment sensitivity to Temozolomide and ionizing radiation in human glioblastoma cells (14), while the loss of primary cilia is correlated with genomic instability in medulloblastoma (48). Kidney tubular cells exhibiting reduced ciliary length demonstrated an increased sensitivity to apoptosis induced by cisplatin, as evidenced in previous studies (49). Similarly, the deficiency of IFT88 in proximal tubular cells, explored through an in vivo knockout (KO) murine model, also resulted in heightened sensitivity to cisplatin (50). Collectively, these observations underscore the pivotal role of primary cilia in the maintenance of genomic integrity and in modulating the cellular response to DNA damage across various cell types and conditions.

The fact that deciliated cells have decreased levels of p53 and p21 and increased levels of γH2AX and Rad51 indicates that their DNA damage response and repair mechanisms are deregulated. However, the ciliary-dependent mechanisms regulating these processes remain unknown. A ciliary pathway that may be implicated is related to Liver kinase B1 (LKB1). Found in both ciliary and cytoplasmic locations, LKB1 plays a role in the DNA damage response by phosphorylating various downstream proteins (51). Its deficiency leads to delayed DNA damage repair (51). Treatment with ATP, which activates LKB1, induces an increase in p-PTEN and inhibits AKT, but only in ciliated cells (42). PTEN, in turn, regulates p53 expression and activity through phosphatase-dependent and expression mechanisms, and also contributes significantly to DNA damage repair and response (52). For example, upregulation of PTEN inhibits AKT and MDM2, which also increase levels of p53 (53). Other pathways that may be involved include Wnt and Hedgehog (54). Activation of β-catenin by Wnt is indirectly involved in activation of p53 and expression of H2AX (55). Hedgehog-signaling GLI proteins can interfere with almost all DNA repair types (56).

The relationship between cilia and DNA repair is particularly relevant for cholangiocytes, as they are exposed to genotoxins generated as hepatic metabolic end products (40). The elevated γH2AX levels observed in Alb-Cre IFT88 mice underscore the potential relevance of our findings to biliary diseases characterized by reduced primary cilia expression in cholangiocytes, such as polycystic liver disease (PLD) (57) and primary sclerosing cholangitis (PSC) (58). Notably, polycystic kidney disease (PKD)—which shares similar mutations with PLD—also exhibits elevated γH2AX levels (59, 60). Moreover, PSC is marked by a decline in primary cilia and an increase in γH2AX, particularly as the disease advances toward a malignant phenotype (58). Collectively, these observations, along with our results, highlight the promise of ciliotherapies as a therapeutic strategy for these ciliopathies. The absence of primary cilia in tumor cells could alter the efficiency of DNA repair processes and be at least one aspect that links the primary cilium with its tumor suppressor characteristics. Providing a potential molecular mechanism may lead to new therapies for CCA, which currently has dismal prognosis with no effective therapies.

## ACKNOWLEDGEMENTS

We would like to thank Tanner Conway and Todd Schuster for expert technical assistance.

## AUTHOR CONTRIBUTIONS

Conceived and designed research: EP, KP, SR, WC, SG; performed experiments: EP, SR; analyzed data: EP, KP, SR, JA; interpreted results of experiments: EP, KP, SR, WC, JA, SG; prepared figures: EP, KP, SG; drafted manuscript: EP; edited and revised manuscript: EP, SG; approved final version of manuscript: EP, KP, SR, JA, WC, SG.

## FIGURE LEGENDS

**Supplementary Figure 1.** (A) Western blot analysis of IFT20 and KIF3A expression in clones NT#4, IFT20#33, KIF3A#1, KIF3A#2, and KIF3A#17. (B) Western blot analysis of IFT88 expression in clones NT#4, IFT88#13, and IFT88#19. NT: non-targeting construct.

**Supplementary Figure 2.** Cell cycle and apoptosis analysis of ciliated and deciliated cells following genotoxic stress. Cells were treated with cisplatin for 24 hours before analysis or exposed to 5 Gy irradiation or 1 mM MMS, followed by a 24-hour incubation before analysis. (A) Representative flow cytometry plots and software adjustments for cell cycle analysis. The FSC/SSC plot was used to exclude small debris, the PI plot to remove doublets, and the DNA content histogram to define cell cycle stages. The histogram shows the distribution of cells across G1 (red), S (striped), and G2/M (solid red) phases. Peaks correspond to diploid G1 and G2 phases, with an intermediate S-phase population (B) Stacked bar graphs representing the percentage of cells in apoptosis, late apoptosis/early necrosis, and necrosis of ciliated and deciliated cells on basal conditions or 24 hrs after 2.5 µM cisplatin, 1 mM MMS or 5 Gy. (C) Representative flow cytometry plots showing FSC/SSC plot to exclude small debris, SSC-H/SSC-A plot to exclude doublets, and Annexin V-FITC/PI dot plot to identify apoptotic cells.

**Supplementary Figure 3.** Immunofluorescence analysis of RAD51 in ciliated and deciliated NHC cells 24 hours after 5 Gy irradiation or in untreated controls. (One-way ANOVA, ****p<0.0001, 630X).

### Declaration of generative AI and AI-assisted technologies in the writing process

During the preparation of this work the author(s) used ChatGPT to check and correct the English language. After using this tool/service, the author(s) reviewed and edited the content as needed and take(s) full responsibility for the content of the publication.

## Bibliography

1. Marzioni M, Invernizzi P, Candelaresi C, Maggioni M, Saccomanno S, Selmi C, Rychlicki C, Agostinelli L, Cassani B, Miozzo M, Pasini S, Fava G, Alpini G, and Benedetti A. Human cholangiocarcinoma development is associated with dysregulation of opioidergic modulation of cholangiocyte growth. Digestive and liver disease: official journal of the Italian Society of Gastroenterology and the Italian Association for the Study of the Liver 41: 523–533, 2009.

2. Banales JM, Cardinale V, Carpino G, Marzioni M, Andersen JB, Invernizzi P, Lind GE, Folseraas T, Forbes SJ, Fouassier L, Geier A, Calvisi DF, Mertens JC, Trauner M, Benedetti A, Maroni L, Vaquero J, Macias RI, Raggi C, Perugorria MJ, Gaudio E, Boberg KM, Marin JJ, and Alvaro D. Expert consensus document: Cholangiocarcinoma: current knowledge and future perspectives consensus statement from the European Network for the Study of Cholangiocarcinoma (ENS-CCA). Nat Rev Gastroenterol Hepatol 13: 261–280, 2016.

3. Rizvi S, and Gores GJ. Emerging molecular therapeutic targets for cholangiocarcinoma. J Hepatol 2017.

4. Ayala D, and Blackstock AW. Effective treatment strategies for cholangiocarcinoma: the challenge remains. Gastrointestinal cancer research: GCR 2: 251–252, 2008.

5. Banales JM, Marin JJG, Lamarca A, Rodrigues PM, Khan SA, Roberts LR, Cardinale V, Carpino G, Andersen JB, Braconi C, Calvisi DF, Perugorria MJ, Fabris L, Boulter L, Macias RIR, Gaudio E, Alvaro D, Gradilone SA, Strazzabosco M, Marzioni M, Coulouarn C, Fouassier L, Raggi C, Invernizzi P, Mertens JC, Moncsek A, Rizvi S, Heimbach J, Koerkamp BG, Bruix J, Forner A, Bridgewater J, Valle JW, and Gores GJ. Cholangiocarcinoma 2020: the next horizon in mechanisms and management. Nat Rev Gastroenterol Hepatol 17: 557–588, 2020.

6. Masyuk AI, Gradilone SA, Banales JM, Huang BQ, Masyuk TV, Lee SO, Splinter PL, Stroope AJ, and LaRusso NF. Cholangiocyte primary cilia are chemosensory organelles that detect biliary nucleotides via P2Y12 purinergic receptors. Am J Physiol Gastrointest Liver Physiol 295: G725–G734, 2008.

7. Masyuk AI, Masyuk TV, and LaRusso NF. Cholangiocyte primary cilia in liver health and disease. Dev Dyn 237: 2007–2012, 2008.

8. Gradilone SA, Radtke BN, Bogert PS, Huang BQ, Gajdos GB, and LaRusso NF. HDAC6 inhibition restores ciliary expression and decreases tumor growth. Cancer Res 73: 2259–2270, 2013.

9. Mansini AP, Lorenzo Pisarello MJ, Thelen KM, Cruz-Reyes M, Peixoto E, Jin S, Howard BN, Trussoni CE, Gajdos GB, LaRusso NF, Perugorria MJ, Banales JM, and Gradilone SA. MiR-433 and miR-22 dysregulations induce HDAC6 overexpression and ciliary loss in cholangiocarcinoma. Hepatology 561–573, 2018.

10. Razumilava N, Gradilone SA, Smoot RL, Mertens JC, Bronk SF, Sirica AE, and Gores GJ. Non-canonical Hedgehog signaling contributes to chemotaxis in cholangiocarcinoma. Journal of hepatology 60: 599–605, 2014.

11. Peixoto E, Jin S, Thelen K, Biswas A, Richard S, Morleo M, Mansini A, Holtorf S, Carbone F, Pastore N, Ballabio A, Franco B, and Gradilone SA. HDAC6-dependent ciliophagy is involved in ciliary loss and cholangiocarcinoma growth in human cells and murine models. Am J Physiol Gastrointest Liver Physiol 318: G1022–G1033, 2020.

12. Pant K, Peixoto E, Richard S, Biswas A, O’Sullivan MG, Giama N, Ha Y, Yin J, Carotenuto P, Salati M, Ren Y, Yang R, Franco B, Roberts LR, and Gradilone SA. Histone Deacetylase Sirtuin 1 Promotes Loss of Primary Cilia in Cholangiocarcinoma. Hepatology 74: 3235–3248, 2021.

13. Chen TY, Huang BM, Tang TK, Chao YY, Xiao XY, Lee PR, Yang LY, and Wang CY. Genotoxic stress-activated DNA-PK-p53 cascade and autophagy cooperatively induce ciliogenesis to maintain the DNA damage response. Cell Death Differ 28: 1865–1879, 2021.

14. Wei L, Ma W, Cai H, Peng SP, Tian HB, Wang JF, Gao L, and He JP. Inhibition of Ciliogenesis Enhances the Cellular Sensitivity to Temozolomide and Ionizing Radiation in Human Glioblastoma Cells. Biomed Environ Sci 35: 419–436, 2022.

15. Johnson CA, and Collis SJ. Ciliogenesis and the DNA damage response: a stressful relationship. Cilia 5: 2016.

16. Fukasawa K, Choi T, Kuriyama R, Rulong S, and Vande Woude GF. Abnormal centrosome amplification in the absence of p53. Science 271: 1744–1747, 1996.

17. Xu X, Weaver Z, Linke SP, Li C, Gotay J, Wang XW, Harris CC, Ried T, and Deng CX. Centrosome amplification and a defective G2-M cell cycle checkpoint induce genetic instability in BRCA1 exon 11 isoform-deficient cells. Mol Cell 3: 389–395, 1999.

18. Sivasubramaniam S, Sun XM, Pan YR, Wang SH, and Lee EYHP. Cep164 is a mediator protein required for the maintenance of genomic stability through modulation of MDC1, RPA, and CHK1. Gene Dev 22: 587–600, 2008.

19. Chaki M, Airik R, Ghosh AK, Giles RH, Chen R, Slaats GG, Wang H, Hurd TW, Zhou W, Cluckey A, Gee HY, Ramaswami G, Hong CJ, Hamilton BA, Cervenka I, Ganji RS, Bryja V, Arts HH, van Reeuwijk J, Oud MM, Letteboer SJ, Roepman R, Husson H, Ibraghimov-Beskrovnaya O, Yasunaga T, Walz G, Eley L, Sayer JA, Schermer B, Liebau MC, Benzing T, Le Corre S, Drummond I, Janssen S, Allen SJ, Natarajan S, O’Toole JF, Attanasio M, Saunier S, Antignac C, Koenekoop RK, Ren H, Lopez I, Nayir A, Stoetzel C, Dollfus H, Massoudi R, Gleeson JG, Andreoli SP, Doherty DG, Lindstrad A, Golzio C, Katsanis N, Pape L, Abboud EB, Al-Rajhi AA, Lewis RA, Omran H, Lee EY, Wang S, Sekiguchi JM, Saunders R, Johnson CA, Garner E, Vanselow K, Andersen JS, Shlomai J, Nurnberg G, Nurnberg P, Levy S, Smogorzewska A, Otto EA, and Hildebrandt F. Exome capture reveals ZNF423 and CEP164 mutations, linking renal ciliopathies to DNA damage response signaling. Cell 150: 533–548, 2012.

20. Pavan ICB, de Oliveira AP, Dias PRF, Basei FL, Issayama LK, Ferezin CD, Silva FR, de Oliveira ALR, Moura LAD, Martins MB, Simabuco FM, and Kobarg J. On Broken Ne(c)ks and Broken DNA: The Role of Human NEKs in the DNA Damage Response. Cells 10: 2021.

21. Choi HJ, Lin JR, Vannier JB, Slaats GG, Kile AC, Paulsen RD, Manning DK, Beier DR, Giles RH, Boulton SJ, and Cimprich KA. NEK8 links the ATR-regulated replication stress response and S phase CDK activity to renal ciliopathies. Mol Cell 51: 423–439, 2013.

22. Baller J, Kono T, Herman A, and Zhang Y. CHURP: A Lightweight CLI Framework to Enable Novice Users to Analyze Sequencing Datasets in Parallel. Pearc ‘19: Proceedings of the Practice and Experience in Advanced Research Computing on Rise of the Machines (Learning) 2019.

23. Joplin R, Strain AJ, and Neuberger JM. Immuno-isolation and culture of biliary epithelial cells from normal human liver. In Vitro Cell Dev Biol 25: 1189–1192, 1989.

24. Follit JA, Tuft RA, Fogarty KE, and Pazour GJ. The intraflagellar transport protein IFT20 is associated with the Golgi complex and is required for cilia assembly. Mol Biol Cell 17: 3781–3792, 2006.

25. Cullen CL, O’Rourke M, Beasley SJ, Auderset L, Zhen Y, Pepper RE, Gasperini R, and Young KM. Kif3a deletion prevents primary cilia assembly on oligodendrocyte progenitor cells, reduces oligodendrogenesis and impairs fine motor function. Glia 69: 1184–1203, 2021.

26. Grentner A, Ragueneau E, Gong C, Prinz A, Gansberger S, Oyarzun I, Hermjakob H, and Griss J. ReactomeGSA: new features to simplify public data reuse. Bioinformatics 40: 2024.

27. Griss J, Viteri G, Sidiropoulos K, Nguyen V, Fabregat A, and Hermjakob H. ReactomeGSA-Efficient Multi-Omics Comparative Pathway Analysis. Molecular & Cellular Proteomics 19: 2020.

28. Lomax ME, Folkes LK, and O’Neill P. Biological consequences of radiation-induced DNA damage: relevance to radiotherapy. Clin Oncol (R Coll Radiol) 25: 578–585, 2013.

29. Pascucci B, Russo MT, Crescenzi M, Bignami M, and Dogliotti E. The accumulation of MMS-induced single strand breaks in G1 phase is recombinogenic in DNA polymerase beta defective mammalian cells. Nucleic Acids Res 33: 280–288, 2005.

30. Duan M, Ulibarri J, Liu KJ, and Mao P. Role of Nucleotide Excision Repair in Cisplatin Resistance. Int J Mol Sci 21: 2020.

31. Scully R, and Xie A. Double strand break repair functions of histone H2AX. Mutat Res 750: 5–14, 2013.

32. Baumann P, and West SC. Role of the human RAD51 protein in homologous recombination and double-stranded-break repair. Trends Biochem Sci 23: 247–251, 1998.

33. Shiojiri N. Analysis of Differentiation of Hepatocytes and Bile Duct Cells in Developing Mouse Liver by Albumin Immunofluorescence: (albumin distribution/liver cells/differentiation/mouse embryos). Dev Growth Differ 26: 555–561, 1984.

34. O’Dell MR, Huang JL, Whitney-Miller CL, Deshpande V, Rothberg P, Grose V, Rossi RM, Zhu AX, Land H, Bardeesy N, and Hezel AF. Kras(G12D) and p53 mutation cause primary intrahepatic cholangiocarcinoma. Cancer Res 72: 1557–1567, 2012.

35. Saha SK, Parachoniak CA, Ghanta KS, Fitamant J, Ross KN, Najem MS, Gurumurthy S, Akbay EA, Sia D, Cornella H, Miltiadous O, Walesky C, Deshpande V, Zhu AX, Hezel AF, Yen KE, Straley KS, Travins J, Popovici-Muller J, Gliser C, Ferrone CR, Apte U, Llovet JM, Wong KK, Ramaswamy S, and Bardeesy N. Mutant IDH inhibits HNF-4alpha to block hepatocyte differentiation and promote biliary cancer. Nature 513: 110–114, 2014.

36. Ikenoue T, Terakado Y, Nakagawa H, Hikiba Y, Fujii T, Matsubara D, Noguchi R, Zhu C, Yamamoto K, Kudo Y, Asaoka Y, Yamaguchi K, Ijichi H, Tateishi K, Fukushima N, Maeda S, Koike K, and Furukawa Y. A novel mouse model of intrahepatic cholangiocarcinoma induced by liver-specific Kras activation and Pten deletion. Sci Rep 6: 23899, 2016.

37. Pant K, Richard S, Peixoto E, Baral S, Yang R, Ren Y, Masyuk TV, LaRusso NF, and Gradilone SA. Cholangiocyte ciliary defects induce sustained epidermal growth factor receptor signaling. Hepatology 2024.

38. Zimmerman KA, Song CJ, Gonzalez-Mize N, Li Z, and Yoder BK. Primary cilia disruption differentially affects the infiltrating and resident macrophage compartment in the liver. Am J Physiol Gastrointest Liver Physiol 314: G677–G689, 2018.

39. Moyer JH, Lee-Tischler MJ, Kwon HY, Schrick JJ, Avner ED, Sweeney WE, Godfrey VL, Cacheiro NL, Wilkinson JE, and Woychik RP. Candidate gene associated with a mutation causing recessive polycystic kidney disease in mice. Science 264: 1329–1333, 1994.

40. Reynolds ES. Environmental aspects of injury and disease: liver and bile ducts. Environ Health Perspect 20: 1–13, 1977.

41. Huang H, Zhu L, Reid BR, Drobny GP, and Hopkins PB. Solution structure of a cisplatin-induced DNA interstrand cross-link. Science 270: 1842–1845, 1995.

42. Mansini AP, Peixoto E, Jin S, Richard S, and Gradilone SA. The chemosensory function of primary cilia regulates cholangiocyte migration, invasion and tumor growth. Hepatology 1582–1598, 2018.

43. Kanai M, Tong WM, Sugihara E, Wang ZQ, Fukasawa K, and Miwa M. Involvement of poly(ADP-Ribose) polymerase 1 and poly(ADP-Ribosyl)ation in regulation of centrosome function. Mol Cell Biol 23: 2451–2462, 2003.

44. Kanai M, Uchida M, Hanai S, Uematsu N, Uchida K, and Miwa M. Poly(ADP-ribose) polymerase localizes to the centrosomes and chromosomes. Biochem Biophys Res Commun 278: 385–389, 2000.

45. Zhang S, Hemmerich P, and Grosse F. Centrosomal localization of DNA damage checkpoint proteins. J Cell Biochem 101: 451–465, 2007.

46. Poruchynsky MS, Komlodi-Pasztor E, Trostel S, Wilkerson J, Regairaz M, Pommier Y, Zhang X, Kumar Maity T, Robey R, Burotto M, Sackett D, Guha U, and Fojo AT. Microtubule-targeting agents augment the toxicity of DNA-damaging agents by disrupting intracellular trafficking of DNA repair proteins. Proc Natl Acad Sci U S A 112: 1571–1576, 2015.

47. Xie H, Wang X, Jin M, Li L, Zhu J, Kang Y, Chen Z, Sun Y, and Zhao C. Cilia regulate meiotic recombination in zebrafish. J Mol Cell Biol 14: 2022.

48. Youn YH, Hou S, Wu CC, Kawauchi D, Orr BA, Robinson GW, Finkelstein D, Taketo MM, Gilbertson RJ, Roussel MF, and Han YG. Primary cilia control translation and the cell cycle in medulloblastoma. Genes Dev 36: 737–751, 2022.

49. Wang S, Wei Q, Dong G, and Dong Z. ERK-mediated suppression of cilia in cisplatin-induced tubular cell apoptosis and acute kidney injury. Biochim Biophys Acta 1832: 1582–1590, 2013.

50. Wang S, Zhuang S, and Dong Z. IFT88 deficiency in proximal tubular cells exaggerates cisplatin-induced injury by suppressing autophagy. Am J Physiol Renal Physiol 321: F269–F277, 2021.

51. Wang YS, Chen J, Cui F, Wang H, Wang S, Hang W, Zeng Q, Quan CS, Zhai YX, Wang JW, Shen XF, Jian YP, Zhao RX, Werle KD, Cui R, Liang J, Li YL, and Xu ZX. LKB1 is a DNA damage response protein that regulates cellular sensitivity to PARP inhibitors. Oncotarget 7: 73389–73401, 2016.

52. Freeman DJ, Li AG, Wei G, Li HH, Kertesz N, Lesche R, Whale AD, Martinez-Diaz H, Rozengurt N, Cardiff RD, Liu X, and Wu H. PTEN tumor suppressor regulates p53 protein levels and activity through phosphatase-dependent and –independent mechanisms. Cancer Cell 3: 117–130, 2003.

53. Ming M, and He YY. PTEN in DNA damage repair. Cancer Lett 319: 125–129, 2012.

54. Shen T, Gao K, Hu Z, Miao Y, and Wang X. Ciliary Mechanism of Regulating Hedgehog and Wnt/beta-Catenin Signaling Modulates Ultraviolet B Irradiation-Induced Photodamage in HaCaT Cells. J Biomed Nanotechnol 15: 196–203, 2019.

55. Karimaian A, Majidinia M, Baghi HB, and Yousefi B. The crosstalk between Wnt/beta-catenin signaling pathway with DNA damage response and oxidative stress: Implications in cancer therapy. DNA Repair 51: 14–19, 2017.

56. Meng E, Hanna A, Samant RS, and Shevde LA. The Impact of Hedgehog Signaling Pathway on DNA Repair Mechanisms in Human Cancer. Cancers (Basel) 7: 1333–1348, 2015.

57. Masyuk AI, Masyuk TV, Trussoni CE, Pirius NE, and LaRusso NF. Autophagy promotes hepatic cystogenesis in polycystic liver disease by depletion of cholangiocyte ciliogenic proteins. Hepatology 75: 1110–1122, 2022.

58. Carpino G, Cardinale V, Folseraas T, Overi D, Grzyb K, Costantini D, Berloco PB, Di Matteo S, Karlsen TH, Alvaro D, and Gaudio E. Neoplastic Transformation of the Peribiliary Stem Cell Niche in Cholangiocarcinoma Arisen in Primary Sclerosing Cholangitis. Hepatology 69: 622–638, 2019.

59. Conduit SE, Davies EM, Ooms LM, Gurung R, McGrath MJ, Hakim S, Cottle DL, Smyth IM, Dyson JM, and Mitchell CA. AKT signaling promotes DNA damage accumulation and proliferation in polycystic kidney disease. Hum Mol Genet 29: 31–48, 2020.

60. Zhang JQJ, Saravanabavan S, Chandra AN, Munt A, Wong ATY, Harris PC, Harris DCH, McKenzie P, Wang Y, and Rangan GK. Up-Regulation of DNA Damage Response Signaling in Autosomal Dominant Polycystic Kidney Disease. Am J Pathol 191: 902–920, 2021.

