## Supplementary figures and images for "Cholangiocytes’ Primary Cilia Regulate DNA Damage Response and Repair"

### Supplementary Figure 1

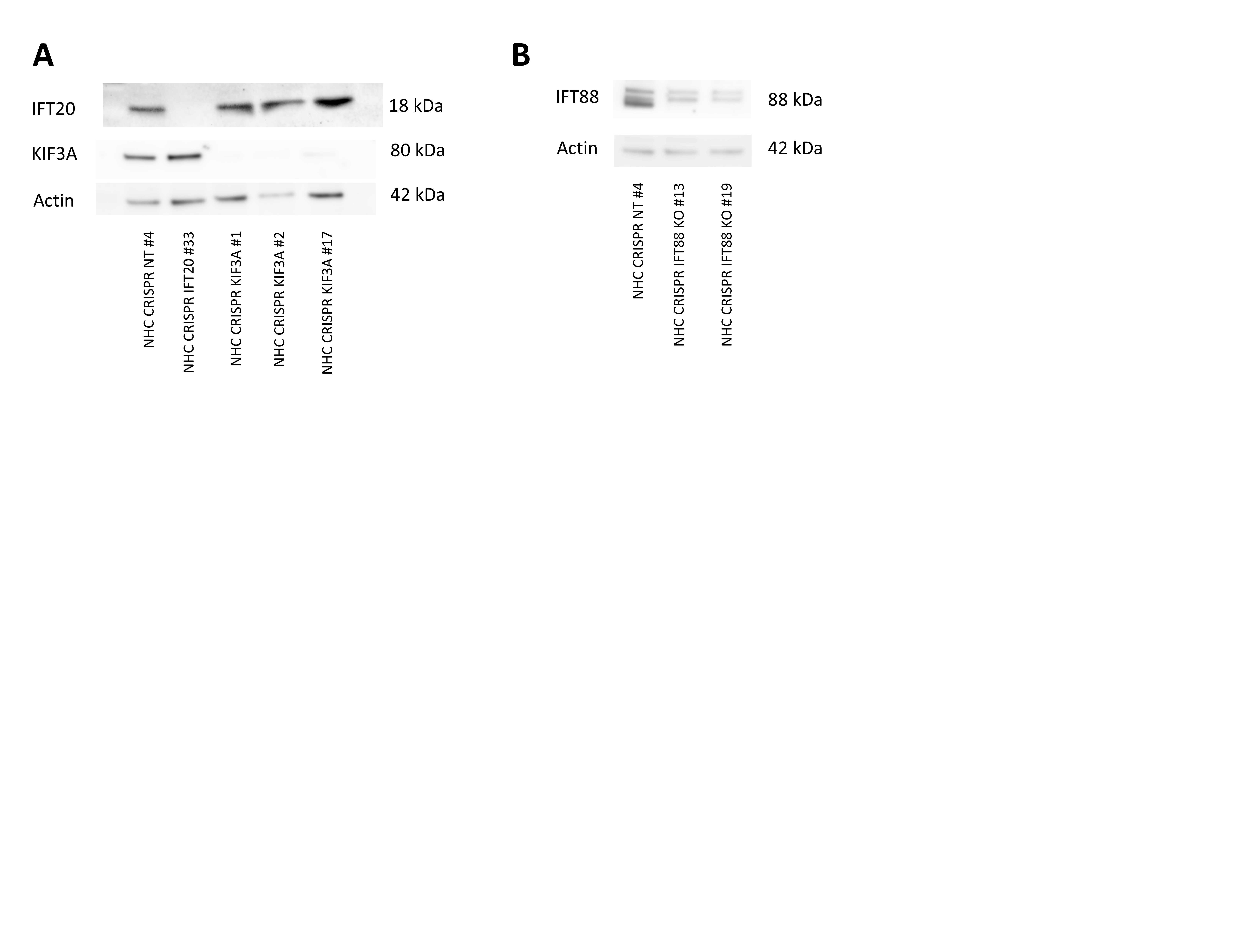

### Supplementary Figure 2

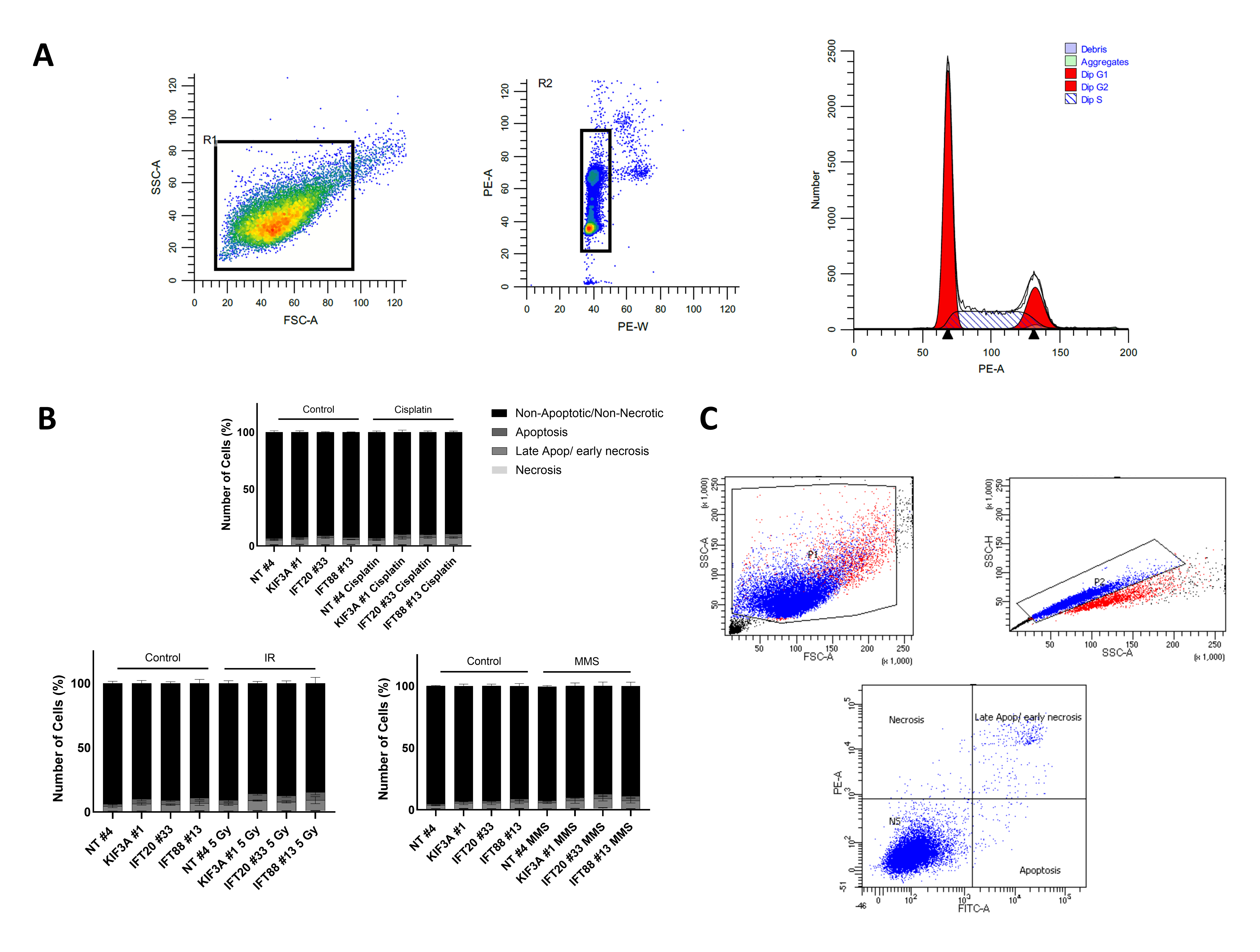

### Supplementary Figure 3

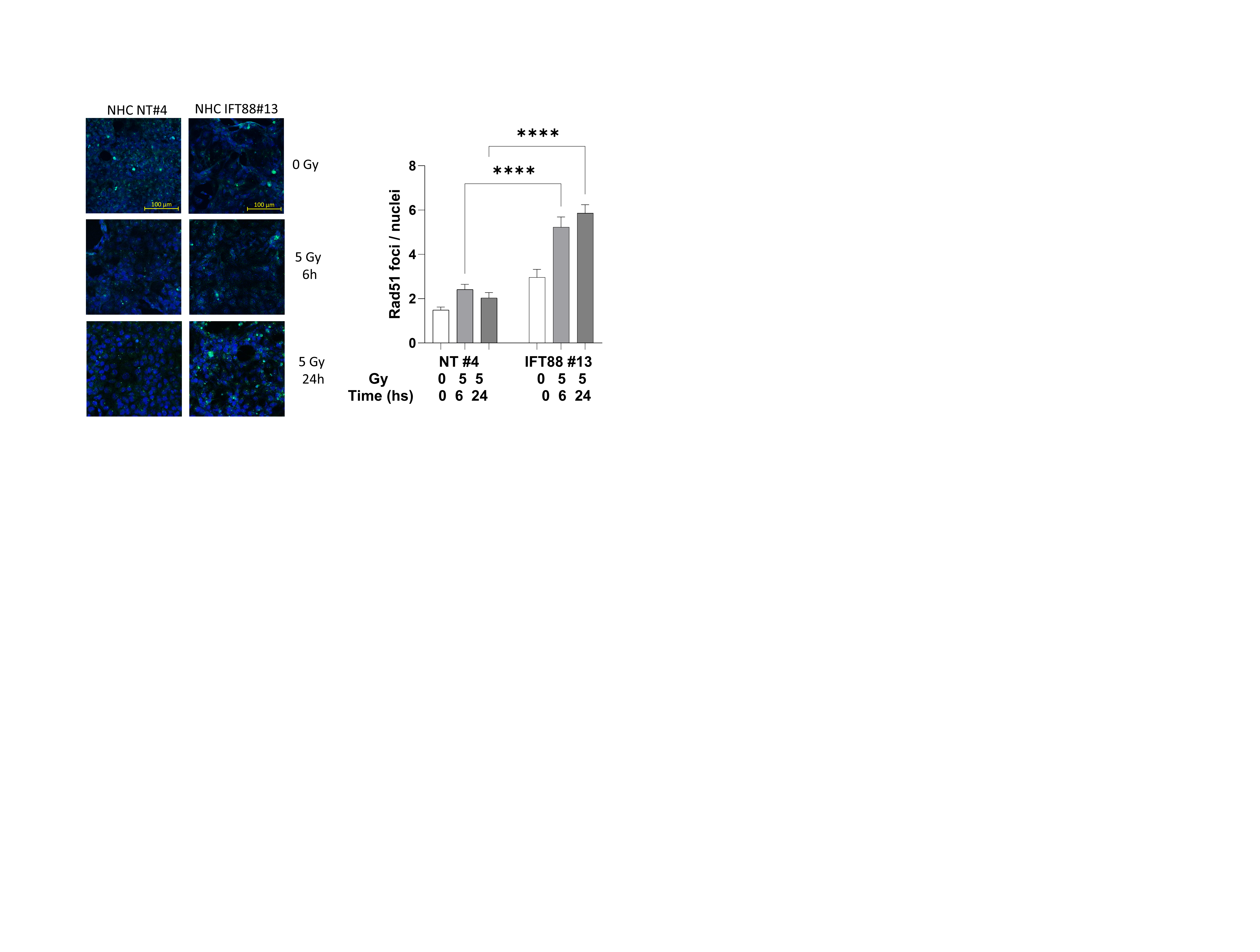
